# Brief exposure of neuronal cells to levels of short chain fatty acids observed in human systemic circulation impair the cell lipid metabolism resulting in associated cell death by apoptosis

**DOI:** 10.1101/2021.06.01.446425

**Authors:** Tiffany A. Fillier, Shrushti Shah, Karen M. Doody, Thu H. Pham, Isabelle Aubry, Michel L. Tremblay, Sukhinder K. Cheema, Jacqueline Blundell, Raymond H. Thomas

## Abstract

Communication between gut microbiota and the brain is an enigma. Alterations in the gut microbial community affects enteric metabolite levels, such as SCFAs. SCFAs have been proposed as a possible mechanism through which the gut microbiome modulate brain health and function. This study analyzed for the first time the effects of SCFAs at levels reported in human systemic circulation on human neuronal cell energy metabolism, viability, survival and the brain lipidome. Cell and rat brain lipidomics was done using UHPLC-HESI-HRAMS/MS. Neuronal cells viability, survival and energy metabolism were analyzed via flow cytometer, immunofluorescence, and SeahorseXF platform. Lipidomics analysis demonstrated that SCFAs significantly remodeled the brain lipidome in vivo and in vitro. The most notable remodulation was observed in the metabolism of phosphatidylethanolamine plasmalogens, and mitochondrial lipids carnitine and cardiolipin. Increased mitochondrial mass, fragmentation, and hyperfusion occurred concomitant with the altered mitochondrial lipid metabolism resulting in decreased neuronal cell respiration, ATP production, and increased cell death. This suggests SCFAs at levels observed in human systemic circulation can adversely alter the brain lipidome and neuronal cell function potentially negatively impacting brain health outcomes.

## Background

The link between enteric (gut) microbiota and the brain in modulating brain health outcome is currently of great interest globally. The gut can communicate with the central nervous system (CNS) through the gut-brain axis, affecting overall brain health. How this communication occurs is not clear ^(6)^. Short chain fatty acids (SCFAs) are proposed as one mechanism through which this communication may occur as they are able to cross the blood-brain barrier ^(6, 30, 40, 48)^. These metabolites are produced by the gut microbiota through bacterial fermentation of fiber in the colon, thus linking them directly to dietary intake of carbohydrates. SCFAs include saturated fatty acids with up to six carbons, but acetate (C2), propionate (C3), and butyrate (C4) in the normal molar ratio of 60:20:20, respectively are the major contributors ^(7, 9, 40)^. Furthermore, upwards of 400-600 mM SCFAs can be produced via dietary fiber fermentation within the colon ^(3)^ and has been shown to rise to 500μM after alcohol dosage ^(7)^. The 60:20:20 ratio reflects the approximate natural occurrence of the molecular ratio of these SCFAs in a healthy human individual, while concentrations reported in circulation of individuals are within the peripheral range of 79±22 μM for total SCFA ^(7)^ and varying peripheral range of 170, 4, and 8 μM for individual SCFAs (acetate, propionate, butyrate respectively) ^(3, 14)^. Within sick individuals, such as in salmonellosis and familial Mediterranean fever (FMF), systemic SCFAs have been observed to be as high as 100uM for propionate alone, with control concentrations being <10uM (25).

SCFAs play a major role in signaling, energy supplementation, homeostasis, and metabolism giving them the ability to elicit cellular, molecular, immunological, and neurological responses ^(9, 23, 30, 40)^. These metabolites have been observed to have beneficial impacts on host health, including their involvement in anti-inflammatory and anti-cancer responses ^(39)^. For example, SCFAs elicit benefits as signaling molecules mediating histone deacetylase (HDAC) inhibitors to modulate various cancers and inflammation ^(21, 23, 39, 40, 48, 53)^. Furthermore, high-fat diets (HFD) enriched with butyrate, and propionate, have been shown to confer obesity resistance, beneficial glucose and lipid metabolism in both mice and humans, versus a non-enriched HFD ^(23, 30)^.

However, in other studies, SCFAs have been implicated in loss of gut integrity, and gut dysbiosis ^(9, 30, 39, 40)^. Gut dysbiosis is an imbalance in the gut microbiome, which affects microbial composition and the metabolites they produce, such as SCFAs. Individuals with dysbiotic and other pathological conditions show on average higher systemic concentrations of SCFAs ranging between 20μM to >1mM ^(25, 30)^. Dysbiosis has been implicated in diseases such as irritable bowel diseases (IBDs) and colorectal cancers (CRC) ^(17)^. Elevated SCFAs have also been reported in the blood circulation of patients with other diseases such as salmonellosis and FMF, as previously mentioned ^(25)^. Environmental factors such as alterations in diet, stress, antibiotic use, or infection can also lead to imbalances in the gut microbial population and elevated SCFAs in the blood of patients ^(6, 23, 48)^. Dysbiotic conditions and the concurrent elevation of SCFAs have been implicated as a potential risk factor in impaired brain health, resulting in alterations in brain cell lipid metabolism and neurodegeneration in Autism ^(28, 41)^, mitochondrial dysfunction, and age-related neurological diseases including Alzheimer’s and Parkinson’s, etc. ^(16, 31, 40, 48)^. Many studies on gut derived SCFA are heavily focused on local or systemic effects within the body. However, there is a lack of research specific to how the brain responds to systemic SCFA concentrations.

The brain lipidome plays several key roles that are both structural and functional in maintaining overall brain health. Diseased or impaired brain function have been associated with changes in brain lipid metabolism, such as those observed in mitochondrial dysfunction in Alzheimer’s and Parkinson’s ^(2, 4, 31, 43)^. Brain mitochondria are critical for maintaining normal brain function and overall brain health since the brain is one of the most energy demanding organs in the body. Mitochondria are the main energy source driving cellular processes within the brain, including neurogenesis, clearance of reactive oxygen species (ROS), neurotransmitter biosynthesis, metabolism and transport of brain lipids, ATP production, and apoptosis susceptibility ^(10, 22, 45, 51)^. Mitochondria dysfunction has been associated with abnormalities in these processes, brain aging and neurodegeneration ^(22, 45)^. SCFAs is known to modulate cellular metabolism and signaling affecting mitochondria function. How SCFAs particularly at concentrations observed in systemic circulation modulate brain lipid metabolism, function and health outcome is poorly understood.

The goal of this study was to determine the potential adverse effects of SCFAs at levels observed in systemic circulation of patience with gut dysbiosis and other pathologic conditions on brain health by analyzing the lipid metabolism in human neuronal cells and Long-Evans rat brain, as well as overall effects on neuronal cell health. We hypothesized that elevated levels of SCFAs, such as those reported in the systemic circulation of patients during gut dysbiosis and other pathologic conditions ^(7, 25)^, can act as a stressor within the brain. Thus, affecting overall brain health via remodeling the brain lipidome, neuronal cell viability, adenosine triphosphate (ATP) production, cell cycle arrest, and possible cell death by apoptosis.

## Materials and Methods

### Reagents

Unless stated otherwise, all reagents were purchased from Sigma-Aldrich (Oakville, ON, CA). SCFA stock (100mM) were prepared in sterile PBS prior to each individual assay.

### In-Vitro Analysis

Cell cultures consisting of SH-SY5Y human neuroblastoma cells were cultured in Dulbecco’s modified Eagle’s medium/Ham’s nutrient mixture F12 (DMEM/F12, 1:1) with 10% fetal bovine serum (FBS) (Fisher Scientific, Ottawa, ON, CA) and 1% Penicillin/Streptomycin (Pen/Strep). Cells were cultured at 37°C and incubated in 5% CO_2_ (High Heat Decontamination CO_2_ Series SCO6AD, Sheldon Manufacturing inc. USA). Prior to lipid analysis, cells were grown in TPP^®^ tissue culture flasks until they reached 80% confluence. Cells were then differentiated to neurons in the presence of 10μM retinoic acid and differentiation media (DMEM/F12 1:1, 3% FBS, 1% Pen/Strep) for 96 hours ^(24)^ in order to represent the most abundant cell type in brain. Differentiation was monitored via microscopy in both control and treated cells. All cells were synchronized in serum-starvation media (containing 0.1% FBS) for 24 hours prior to treatment with SCFAs. Cells were treated with 50μM, 100μM, or 1000μM SCFAs at a 60:20:20 ratio (acetate: propionate: butyrate) for 72 hours. Phosphate buffered saline (PBS) was used as a vehicle control. Prior to flow cytometry, Cytation imaging, and STED imaging, cells were plated in 6-well plates (Corning^®^, VWR International, QC, CA) at a density of 1×10^5^, differentiated for 96 hours, starved for 24 hours, and then treated with 50μM, 100μM, or 1000μM SCFAs (acetate: propionate: butyrate, 60:20:20) for 72 hours. After the 72-hour treatment for all analysis, adherent cells were collected via trypsin-EDTA solution, then pelleted by centrifuging (1500rpm, 5 min), representing a population of approximately 1×10^6^ cells for lipid analysis. Max cell passages did not exceed 20 passages.

### In-Vivo Analysis

Animal study procedures were approved by Memorial University of Newfoundland Animal Care Committee and followed the guidelines of the Canadian Council on Animal Care (protocol number: 16-01-RT). Briefly, 45-50 day old Long-Evans rats from Charles River Laboratories (Canada) were used for this study. A total of 12 males and 12 females were kept under standard conditions and divided into treatments groups (n=6). The control group received 0.1M PBS at 2mg/kg body weight. Treatment groups were given the same mixture and ratio of SCFAs (acetate: propionate: butyrate, 60:20:20, PBS was the vehicle) as compared to the SH-SY5Y cell methods. Rats received 500mg/kg body weight of SCFA via intraperitoneal (IP) injection for 7 consecutive days. IP was chosen to negate the gut metabolism of SCFA and increase the levels reaching the brain in order to directly mimic potential exposure arising from the central circulation system during normal and stress conditions ^(36)^. Long-Evans rat brain samples were collected 20 minutes after the final IP injection, snap-frozen via liquid nitrogen, and stored at −80 until lipid analysis ^(35)^.

### Lipid Analysis

Lipids were extracted from the neuronal cell pellets and homogenized rat brain using the Bligh and Dyer method ^(1)^. The lipids extracted were analyzed using hydrophilic interaction liquid chromatography (HILIC) for acylcarnitine analysis and C30 reverse-phase liquid chromatography (C30RPLC) for lipidome profile by liquid chromatography coupled with high resolution tandem mass spectrometry. The HILIC column was obtained from Phenomenex (Torrance, CA, USA) and the C30RPLC column used was the Accucore C30 column from ThermoFisher Scientific (ON, Canada). HILIC and RPLC methods were derived from previously published methods ^(33)^. Organic acid analysis was also conducted on the lipid extracts in order to analyze SCFAs, 20μL of extract or standard solution was added to 500μL of acetonitrile solution containing 1mM DPDS, 1mM TPP, and 1mM HQ. The reaction mixture was incubated at 60 °C for 60 min. The HQ derivatives in the reaction mixture were analyzed by LC-MS. The LC-MS column used was a Phenomenex C18 Reverse Phase Column (Torrance, CA, USA) in a LTQ Orbitrap. Solvent A consisted of water, 0.05% acetic acid, and 5 mM ammonium acetate, and solvent B consisted of acetonitrile, 0.05% acetic acid, and 5 mM ammonium acetate, with an LC-MS gradient of 5% B to 95% B over a 16-min method and flow rate of 0.3mL/min based on previously published methods ^(27)^.

### Flow Cytometry Analysis

In order to analyze cell cycle progression following neuronal differentiation and incubation with SCFA, acoustic flow cytometry was used to analyze population and cell viability for seeding and cells were treated as stated above. The supernatants were collected, combined with trypsinized adherent cells, and centrifuged to obtain cells as pellets. The cell pellet was fixed with ice cold 80% ethanol for 30 minutes and stored in the fridge overnight, then stained with 20 μg/ml of propidium iodide (PI) in 0.1% Triton x-100 (in PBS) containing 200 μg/ml DNAse-free RNAse. Prior to analysis, cells were vortexed and stored at room temperature (RT) in the dark for 30 minutes. Data were acquired using an Attune Acoustic Focusing Flow Cytometer (Applied Biosystems, ThermoFisher) equipped with Attune Cytometric Software, which was used for both data acquisition and analysis.

### Cellular Imaging using Cytation 3 Imager

All staining and mounting reagents were purchased from Fisher Scientific unless stated otherwise. In order to analyze morphological changes, cells were differentiated to neurons, starved, and treated on coverslips, then stained with fluorophores to discern the nuclei, actin, and mitochondria. For fluorescence staining, cells were first stained with 500nM MitoTracker™ Deep Red (ThermoFisher Scientific,ON, Canada) in 2ml of culture media and incubated at 37°C in 5% CO_2_ for 30 minutes. This was followed by fixing the cells in 4% Paraformaldehyde (PFA) dissolved in PBS. The fixed cells were then permeabilized in 0.2% Triton X-100 in PBS and blocked with 2% FBS in PBS before staining with Phallodin-AF568 (ThermoFisher Scientific,ON, Canada) and Hoechst (ThermoFisher Scientific,ON, Canada). Coverslips containing the stained cells were mounted to slides with ProLong™ Gold Anti-Fade (ThermoFisher Scientific,ON, Canada) mounting media before imaging. Imaging and analysis were performed on a Cytation 3 Cell Imaging Multi-Mode Reader (Biotek, Winooski, VT, U.S.A). Data acquisition and processing was accomplished with Gen 5 software package (Biotek).

### Super Resolution Imaging using STED Microscopy

Differentiated, starved, and treated cells were fixed, permeabilized, and blocked in the same format as above under *Cytation 3 Imaging*. Cells were then initially stained with microtubule-associated protein 2 (MAP2) monoclonal primary antibody (M13) (ThermoFisher Scientific,ON, Canada) at a 1:300 dilution then incubated overnight at 4°C. Following this, staining with Star Red (KK114) (20mins, RT, covered) (Abberior, Göttingen, Germany) and Star 580 (20mins, RT, covered) (Abberior, Göttingen, Germany) was completed. Cells were then analyzed under stimulated emission depletion (STED) microscopy with corresponding confocal microscopy ^(50)^.

### Assessment of Mitochondrial Function using the Seahorse Platform

The effects of SCFA treatment on mitochondrial oxygen consumption rate (OCR) of differentiated SH-SY5Y cells were assessed using a Seahorse XFe96 analyzer (Agilent, Santa Clara, CA, U.S.A). In this assay, additions of the ATP synthase inhibitor oligomycin (2μM), the mitochondrial uncoupler carbonyl cyanide 4-(trifluoromethoxy)phenylhydrazone (FCCP) (1μM), and the complex I inhibitor rotenone (1μM) into the assay medium was used to provide insight into different aspects of mitochondrial function. SH-SY5Y cells were plated at a density of 5000 cells per well in the Seahorse XF96 cell culture micro-plate and differentiated, starved, and treated as explained above in *cell culture*. The Seahorse measurements were carried out as follows: The cell culture media was replaced by the XF (extracellular flux) assay medium supplemented with 10mM Glucose (ThermoFisher, ON, Canada), 2mM Glutamax (Gibco) and 1mM pyruvate (HyClone) and incubated for 1 hour at 37°C and 5% CO_2_ prior to Seahorse measurements. Data acquisition included ATP production, basal respiration, maximal respiration, and spare respiratory capacity. Data was normalized on cell count using the CyQuant proliferation kit (ThermoFisher, ON, Canada), then exported for statistical analysis ^(12)^.

### Statistical Analysis

All statistical analysis was completed using XLSTAT Premium Version (Addinsoft, New York, USA) as previously published ^(46)^. All data was analyzed for normality and outliers. A multivariate approach was applied to the lipid classes determined via Lipid Search 4.2 (ThermoFisher Scientific,ON, Canada) and Xcalibur 3.1 (ThermoFisher Scientific,ON, Canada). Principle components analysis (PCA) was conducted in order to determine if the SCFA treatments were segregated within the same quadrants of the biplot based on lipid alterations. One-way analysis of variance (ANOVA) was used to analyze the lipid molecular species that accounted for the segregation of the treatments and control in different quadrants of the PCA biplots. Fisher’s Least Significant Difference (LSD) was used to separate the means when the treatments were observed to significantly differed from each other at p>0.05. Differential expression was also used to analyze the lipid molecular species to discern differences between the treatments and control. Figures were prepared using XLSTAT (Premium version, Adinsoft, Paris France) and Adobe Illustrator (Adobe, California, USA).

## Results

### SCFA treatment alters global and mitochondrial lipid composition of SH-SY5Y cells and Long-Evans rat brain

In order to determine whether elevated levels of SCFAs observed in central circulation during gut dysbiosis or other pathological conditions modulated brain lipid metabolism and function, two models were used: retinoic-acid differentiated SH-SY5Y human cell line (Fig. 1) and Long Evans rat brain. We observed that exposure of neuronal cells to elevated levels of SCFAs altered the neuronal cell lipidome relative to the control treated cells (Fig. 1D). Most notably, increases in ceramides (Cer & CerG1), lyso-phosphatidylethanolamine (LPE), cardiolipin (CL), carnitine (Car/C0) and acylcarnitines (AC) occurred concomitantly with decreases in phosphatidylcholine (PC). The most significant (p<0.05) alterations occurred in neuronal cells exposed to the 50μM and 1000μM treatments (Fig. 1D). Organic acids, including SCFAs, were analyzed in the brain of Long Evans rats in both control and treated animals. All SCFAs were observed to be increased across male treated samples, with propionic acid showing an increase across female treated samples (Fig. 1E).

**Figure 1.**
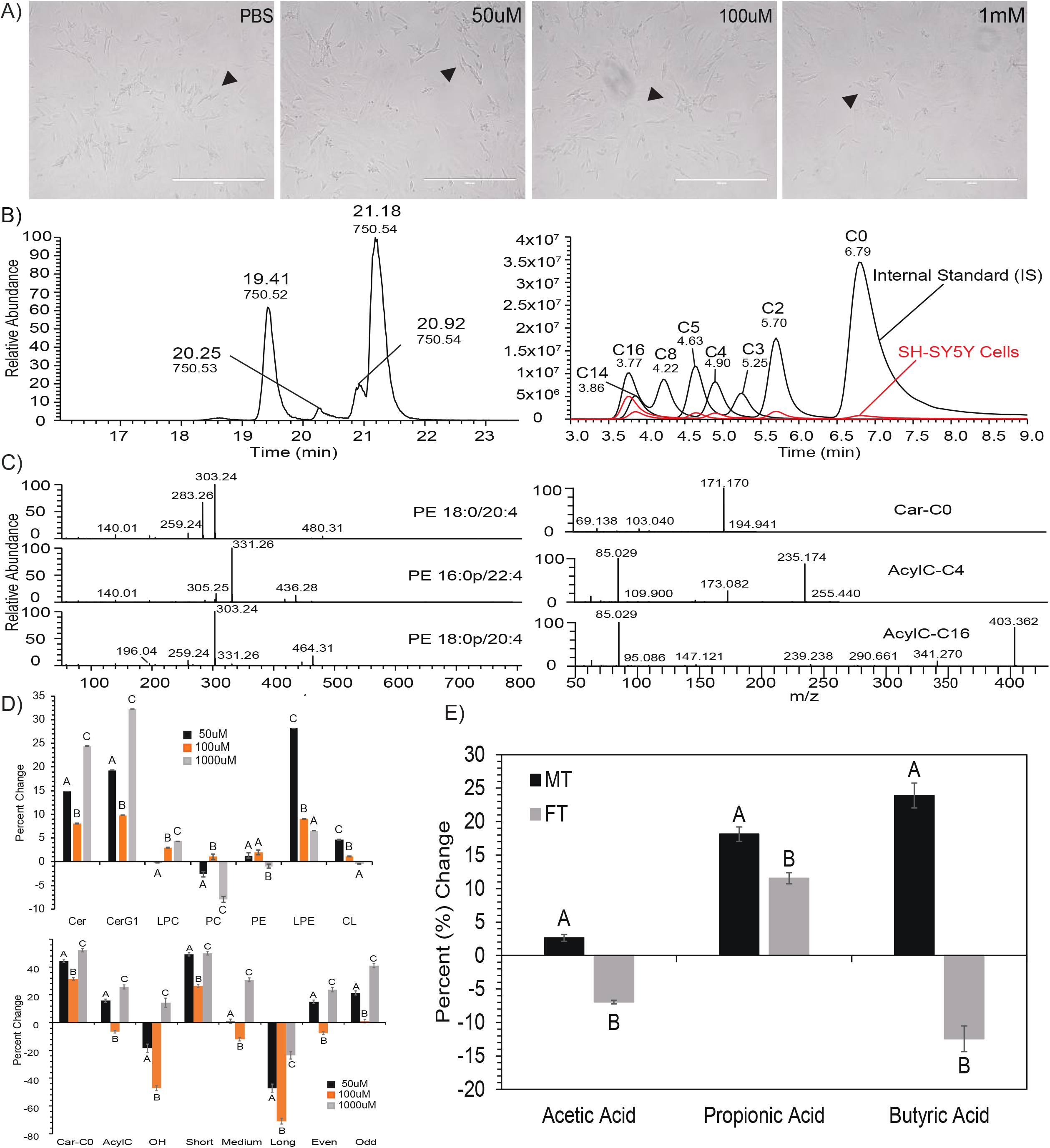
Lipid analysis completed on SH-SY5Y neuroblastoma cells show global alterations due to SCFA treatment. A) Quality of differentiated control and differentiated treated cells. B) C30RPLC and HILIC extraction ion chromatography example (XIC) in −/+ mode respectively of PE (m/z 750) and AC lipids found in cells. C) C30RP spectrum identification of PE18:0/20:4, isomers PE16:0p/22:4, PE18:0p/20:4 and HILIC spectrum identification D-labeled carnitine species Car-C0-D9 (free C), AcylC-C4-D3, and AcylC-C16-D3. D) Percent change by lipid class of treated cells vs control for sphingolipid, phospholipid, and carnitine. E) Alterations in SCFAs in complementory rat study, represented by percent (%) change. Bar charts representative of means ± standard error. Means represented by different superscripts are significantly different at p<0.05. N = 4 samples per experimental treatment concentration (50uM, 100uM, 1000uM) replicate.

Interestingly, some of the most dramatic changes were observed following evaluation of AC. Both AC and CL are lipids specific to the mitochondria. We further evaluated these mitochondrial lipids including their molecular species to better understand how SCFA treatments affected neuronal mitochondrial lipid metabolism. Following PCA analysis of Car/AC, we observed that each of the SCFA treatments clustered in different quadrants of the biplots compared to the control, and this grouping accounted for 83.70% of the total variation present in the AC data (Fig. 2A). The long chain AC clustered with the control in quadrant 3, while the short chain AC clustered with the 50μM and 1000μM SCFA treatments in quadrants 1 and 2 of the PCA biplot (Fig. 2A). Consistent with these groupings, elevated levels of free carnitine and AC (Fig. 2B-C) was observed in neuronal cells treated with all three concentrations of SCFAs, while the level of propionylcarnitine was elevated in both 50μM and 1000μM treatments compared to the control. Furthermore, the levels of all short chain AC including butyl and pentylcarnitine were significantly (p<0.05) elevated in neuronal cells treated with 1000μM SCFAs. Conversely, a general reduction in long chain AC was observed in neurons treated with all three concentrations of SCFAs (Fig 2D). Specifically, cells treated with 50μM and 100μM SCFAs had significantly (p<0.05) lower levels of C14:0, C16:0, C18:0, C18:1, and C20:1 AC compared to the control.

**Figure 2.**
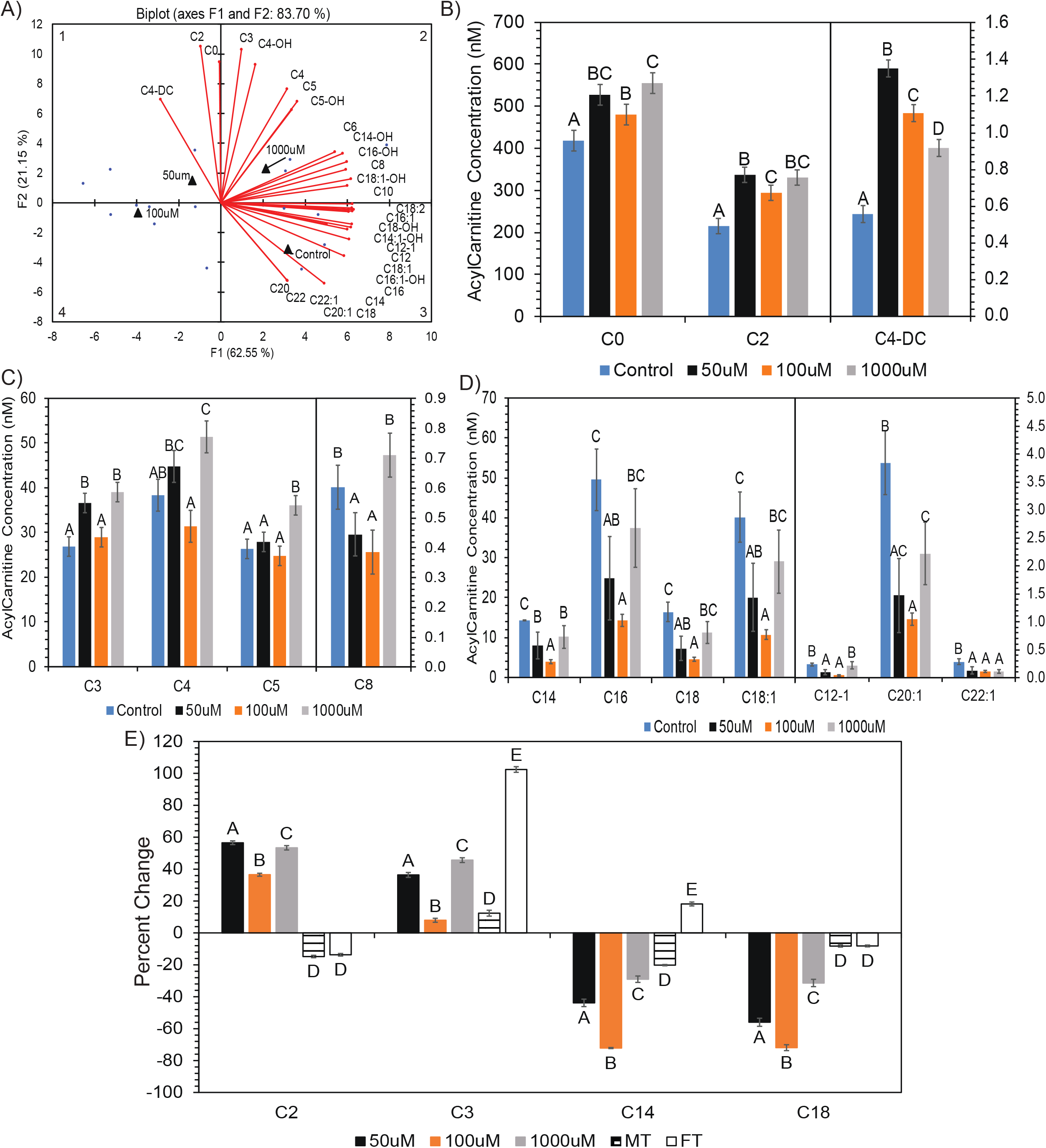
Alteration in carnitine and acyl-carnitine compostition after SCFA treatment. A) PCA biplot showing relationship between SCFA treatment and carnitine molecular species. ANOVA revealed significantly (p<0.05) altered carnitine molecular species in quadrant 1 (B), quadrant 2 (C), quadrant 3 (D) of the PCA. E) Percent change of treatments vs respective controls for ANOVA significant (p<0.05) acyl-carnitine species altered in both SH-SY5Y cell line and Long Evans rat brain. Species determined via PCA and Differential Expression, repspectively, prior to ANOVA. Bar charts representitive of means ± standard error. Means represented by different superscripts are significantly different at p<0.05. In Long Evans rats, N = 6 per experimental replicate. MT = male rats treated with SCFAs. FT = female rats treated with SCFAs.

Based on these findings, we compared the SH-SY5Y cell line AC composition to the Long-Evans rat brain AC composition to determine if the same acylcarnitines were altered in-vitro and in-vivo (Fig. 2E). We observed that C2, C3, C14:0, and C18:0 was altered simultaneously in both cell line and rat brain. The trend between sample sets is different regarding C2, where the cell line is showing an increase and rat brain is showing a decrease in C2. However, C3, C14:0, and C18:0 showed similar increasing (C3) or decreasing (C14:0 & C18:0) trends between the cell line and rat brain samples exposed to SCFAs. The increase in C3 and decrease in C18:0 is observed in the cell line as well as both male treated (MT) and female treated (FT) rats. The decrease in C14:0 is observed in the cell line and only within MT rats.

Similar to the AC, we observed CL clustered in distinct quadrants of the PCA biplot with SCFA treatments (Fig. 3A). Neurons treated with 100μM SCFAs and the control clustered together in quadrant 2 with C16:1 enriched CL molecular species. The 1000μM treated cells clustered in quadrant 4 with predominantly CL molecular species enriched with saturated (C16:0, 18:0) and unsaturated (C18:1) fatty acids; while the 50μM treated cells clustered in quadrant 1 based on a combination of C18:1 and C16:1 enriched molecular species. These groupings accounted for 61.61% of the total variation present in the CL data (Fig. 3A). Furthermore, the CL molecular species that clustered in quadrant 2 were significantly lower in cells treated with 1000μM SCFA compared with the control and other treatments (Fig. 3B), while the molecular species in the cells treated with 50μM and 1000μM SCFAs and clustered in quadrant 4 were significantly higher than that of the control (Fig. 3C). Furthermore, we compared the CL composition in the cell line to the CL composition in the rat brain samples (Fig. 3D-E). Although no altered species between the sample sets had the same fatty acid composition (Supplemental Data, Fig. 1), when comparing total CL alteration, the same increasing trend can be seen in-vitro and in-vivo. Within the cell line, an increase in CL is observed after treatment with 50μM and 1000μM SCFAs. Similarly, within rat brain, an increase in CL was observed in both MT and FT rats compared to controls. However, only the MT rats are significantly different (p>0.05) when compared to respective control.

**Figure 3.**
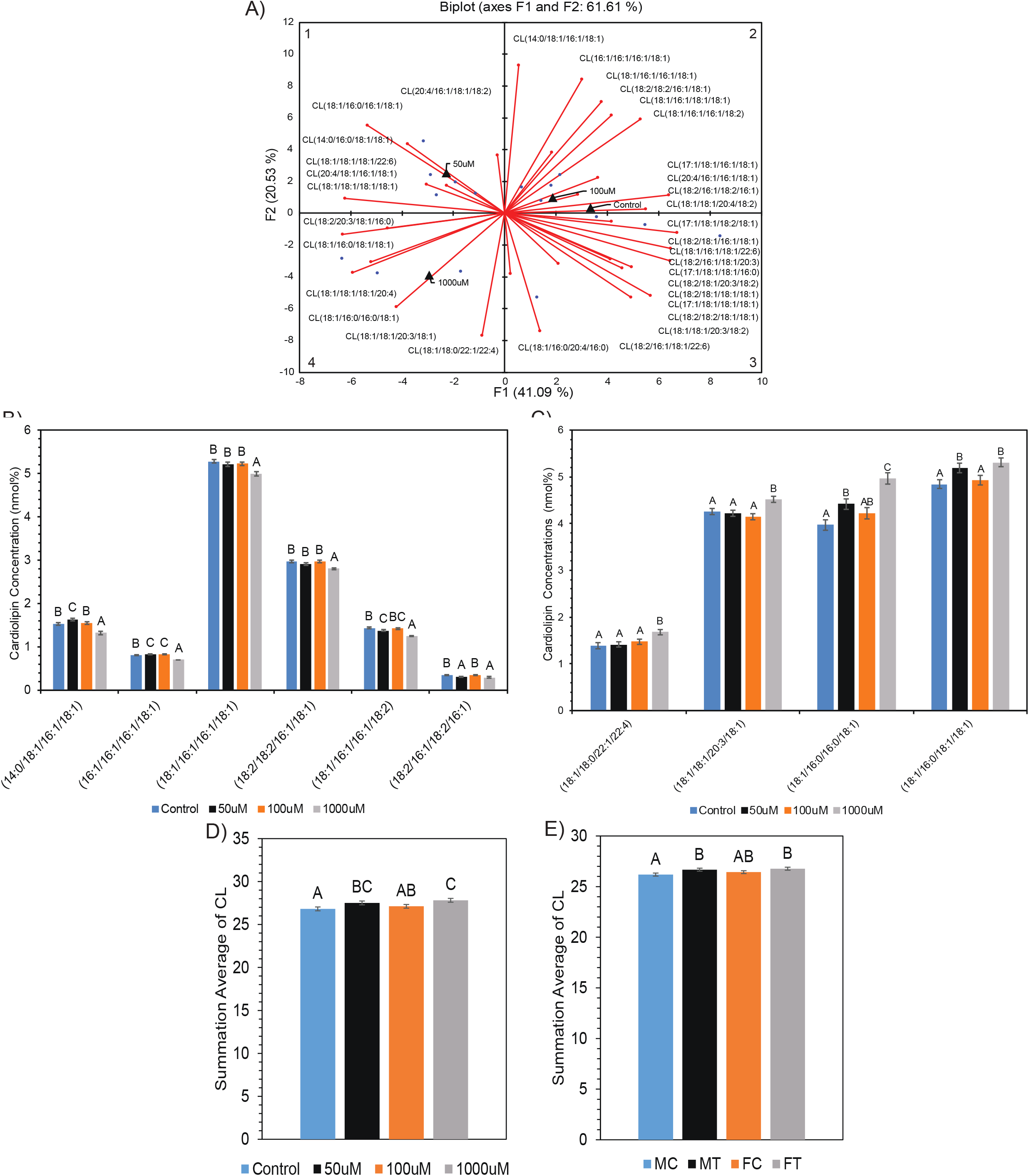
Alteration in cardiolipin (CL) compostition after SCFA treatment. A) PCA biplot showing relationship between SCFA treatment and cardiolipin molecular species. ANOVA revealed significantly (p<0.05) altered CL molecular species in quadrant 2 (B), quadrant 4 (C) of the PCA. Average sum of all ANOVA significant (p<0.05) CL species altered in D) SH-SY5Y cell line and E) Long Evans rat brain. Species determined by PCA and Differential Expression, respectively, prior to ANOVA. Bar charts representitive of means ± standard error. Means represented by different superscripts are significantly different at p<0.05. In Long Evans rats, MC = male control rats, FC = female control rats.

### Treatment with SCFAs alter mitochondrial morphology in SH-SY5Y cells

Following the observation that SCFAs significantly altered the composition of the mitochondrial lipids AC and CL, we determined whether exposure of neuronal cells to SCFAs also affected mitochondrial structure and morphology using MitoTracker™ Deep Red staining followed by cytation and super resolution imaging. The results from the cytation image analysis revealed an increase in functional mitochondrial mass (as seen by increase in average MitoTracker™ intensity) in SCFA treated SH-SY5Y cells (p<0.05) (Fig. 4A-B). Super-resolution (STED) imaging was then used to better assess the mitochondrial morphology in greater detail. This analysis showed normal mitochondrial morphology in both the control and 100μM SCFA treatments, which corresponds to trends observed with the CL lipid analysis. However, STED analysis of the 50μM and 1000μM treatment showed a high proportion of fragmented and hyperfused mitochondria, respectively (Figure 4C-D).

**Figure 4.**
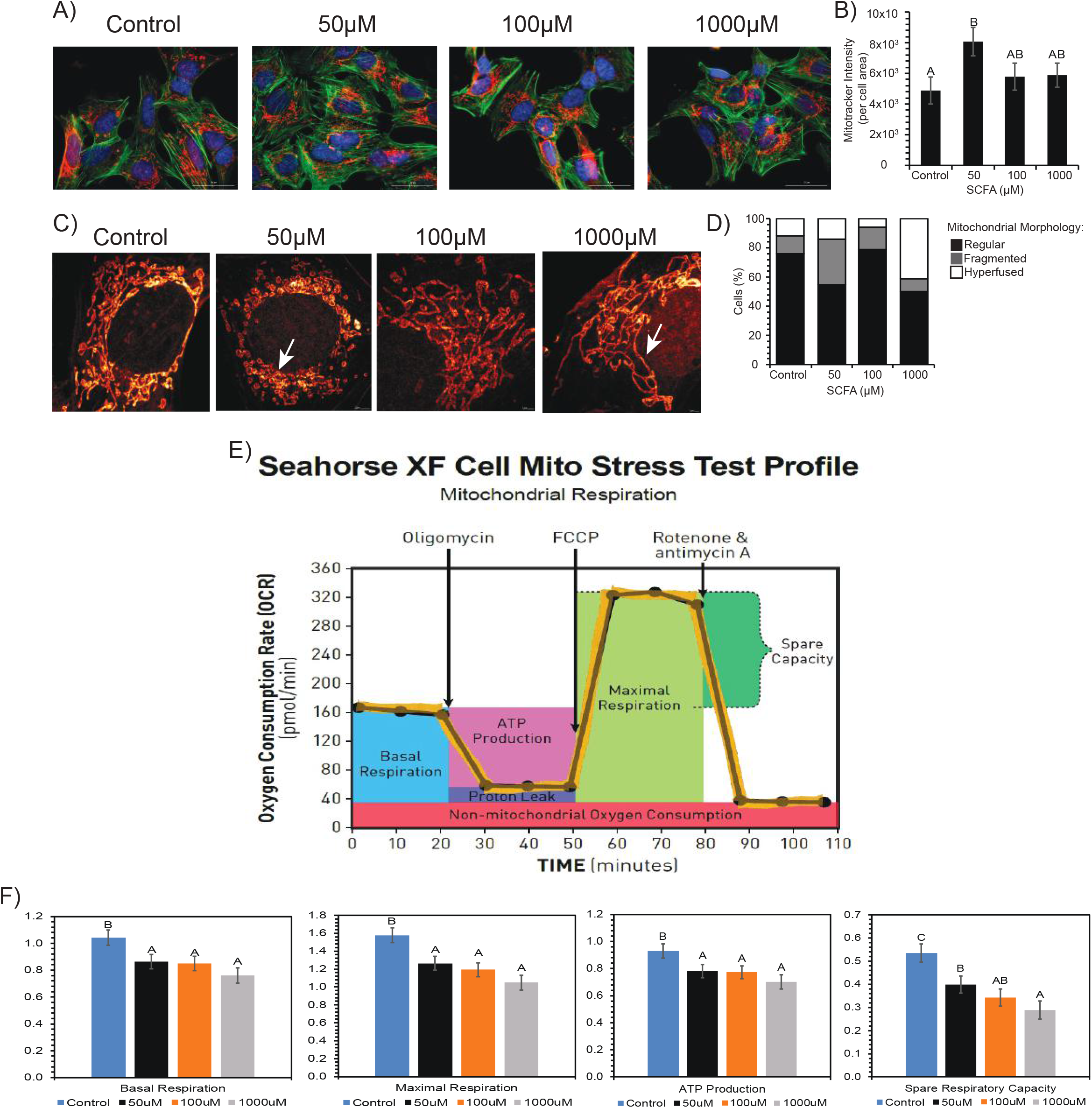
Morphological changes observed in the mitochondria between control vs SCFA treated cells. A) Mitotracker Deep Red under Cytation3 Imaging revealed increased dye intensity after treatment. B) Bar graph numerically represents increase in dye intensity. Bars represent mean ± standard error. Different superscripts represent statistically significant increase (p<0.05). C) STED imaging revealed fragmented (50uM) and hyperfused (1000uM) mitochondria after treatment. D) 100% stacked column graph numerically represents relationship of regular, fragmented and hyperfused mitochondria per treatment. (E,F) Decrease in respiratory capacity and ATP production in cells after treatment.E) Representative of general mitochondrial stress test profile and outputs from Seahorse XF as it measures OCR over time (from Agilent). F) Seahorse XF analysis results after SCFA treatment showing an overall decrease in basal respiration,maximal respiration, ATP production, and spare respiratory capacity. Bars represent means ± standard error. Different superscripts represent statistically different (p<0.05) decreases in respiration and ATP.

### Decreased respiratory ability and ATP production in SH-SY5Y cells after treatment with SCFAs

Considering increased mitochondrial mass and abnormal mitochondrial morphologies were observed in SH-SY5Y cells upon SCFA treatment, we next determined the effect of SCFAs on mitochondrial function as determined by cellular respiration, ATP production, and mitochondrial stress using the Seahorse XF Cell Mito Stress Test platform (Fig. 4E). Significant decreases (p<0.05) in OCR (oxygen consumption rate) measurements; basal respiration, maximal respiration, ATP production, and spare respiratory capacity (SRC), were observed in a dose response manner. Consistently, we observed that ATP production, SRC, basal and maximal respiration significantly decreased (p<0.007) across all treatment concentrations, indicating SCFA treatments suppressed mitochondrial respiration and ATP production in neuronal cells (Fig. 4F).

### SCFAs alter ceramide and plasmalogen composition in-vitro and in-vivo, concurrent with impaired neuronal cell division and induced cell death in SH-SY5Y cells

Given the mitochondrial stress observed in SH-SY5Y cells upon SCFA exposure, we next determined whether the composition of ceramide and plasmalogen lipids were altered in-vitro and in-vivo in response to SCFA treatment. Ceramide and plasmalogen can be used as markers for mitochondrial stress. Although increases and decreases occur in both lipid classes, depending on molecular species (Supplemental Data, Fig. 2), a significant increase in the ceramide:plasmalogen ratio in neuronal cells treated with 50μM and 1000μM SCFA was observed (Fig. 5A). PCA analysis further confirmed, via quadrant segregation, the greatest variations in plasmalogens and ceramides occurred within the 50μM and 1000μM treatments (Fig. 5B). Given the vast but various alteration in molecular species, we analyzed overall trend and fold-change. We determined that the increase in the ceramide:plasmalogen ratio was due to an overall increase in both ceramide and plasmalogen concentrations after 1000μM SCFA treatment (Fig. 5C-D). However, the fold-change for ceramides (1.32) was higher than that of plasmalogens (1.14) by approximately 15%, thus the increased ratio was mostly due to increased ceramide concentration.

**Figure 5.**
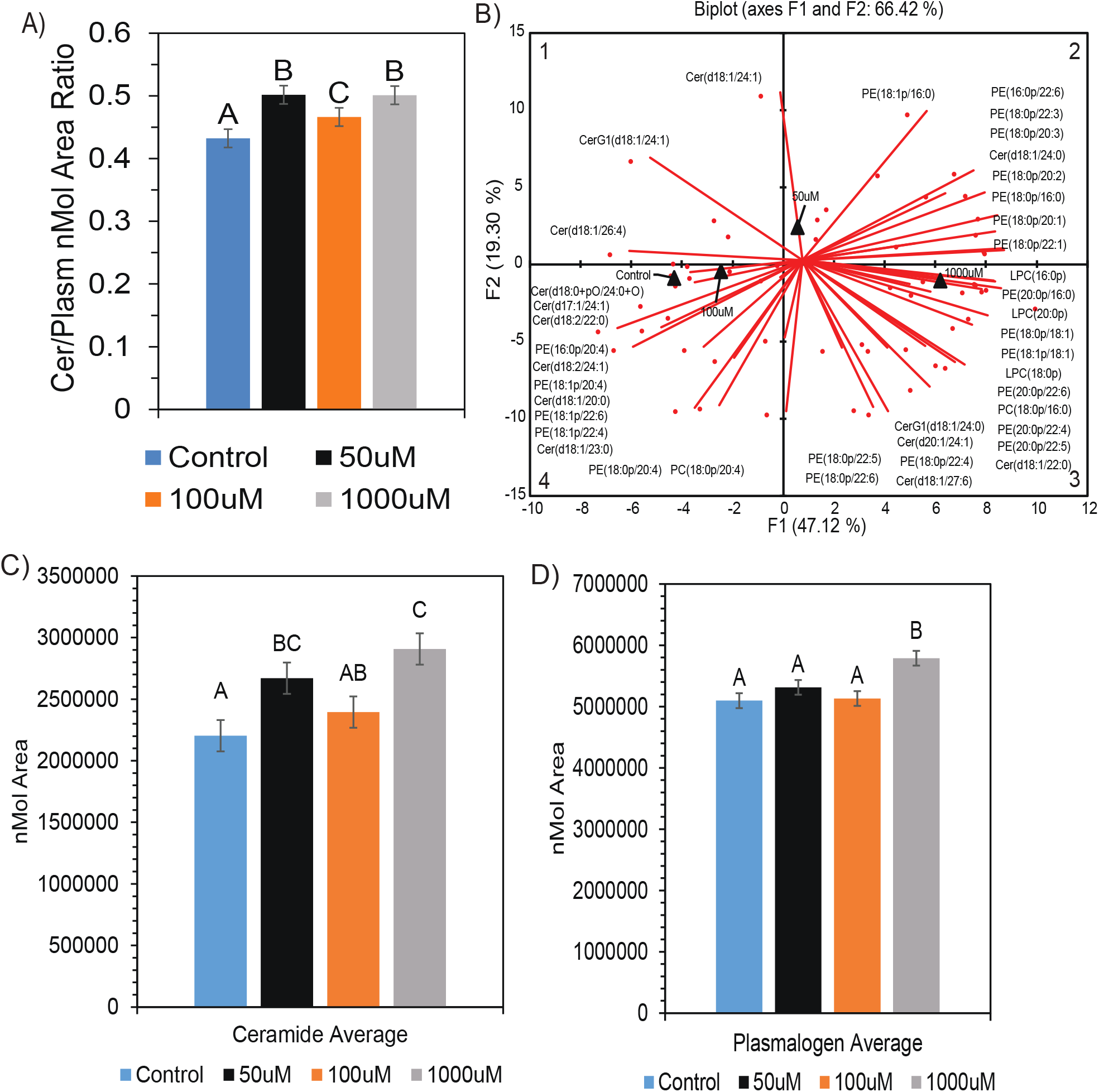
Alteration in ceramide and plasmalogen composition in neuronal cells after SCFA treatment. A) Increased ratio of ceramide:plasmalogens after treatment. B) PCA biplot showing alteration in both ceramide and plasmalogen molecular species between control/100μM, 50μM, and 1000μM SCFA treatment. ANOVA of average nmol area for both ceramide (C) and plasmalogen (D) showed significant (p<0.05) difference after SCFA treatment. Ceramides show an average fold-change increase of 1.32 and plasmalogens show an average fold-change increase of 1.14 after 1000μM treatment. Bar charts representitive of means ± standard error. Means represented by different superscripts are significantly different at p<0.05.

Furthermore, we determined whether the alteration in ceramides and plasmalogens observed in SH-SY5Y cell line were the same in Long-Evans rat brain (Fig. 6A). The observed increase of ceramide and plasmalogen concentrations were sustained in analysis of ceramide and plasmalogen molecular species. This trend was observed in 50μM and 1000μM treated cells and female treated rats in respect to phosphatidylethanolamine (PE) (18:0p/16:0), PE(18:0p/20:3), PE(16:0/22:6), PE(20:0p/22:4), and Cer(18:1/24:0). PE(18:0p/22:4) increased in both 1000μM treated cells and female treated rats. We observed a decreased in plasmalogen species enriched C20:4 fatty acid in both neuronal cells and female treated rat brain following treatment with SCFAs. Female treated rats show the most similarity to the 1000μM SCFA treatment. The increased ceramide:plasmalogen ratio reflects a potentially pro-apoptotic shift in lipid composition in SH-SY5Y cells upon treatment with SCFA, and so we next determined the effect of SCFA treatment on cell cycle progression. Cell cycle analysis of SH-SY5Y cells treated with SCFA revealed that there was a significant decrease (p<0.0001) in cells in the G_2_/M phase (Fig. 6B,D) across all treatment concentrations compared to control cells, while a significant increase (p<0.0001) in cells was observed in the Sub-G_0_ phase (apoptotic cells) (Fig. 6B,C). These results suggest that SCFA elicit a dramatic reduction in neurons in the G2/M phase and that this is accompanied by induction of cell death, represented by accumulation of cells in the sub-G0 region of the cell cycle (Fig. 6B-D).

**Figure 6.**
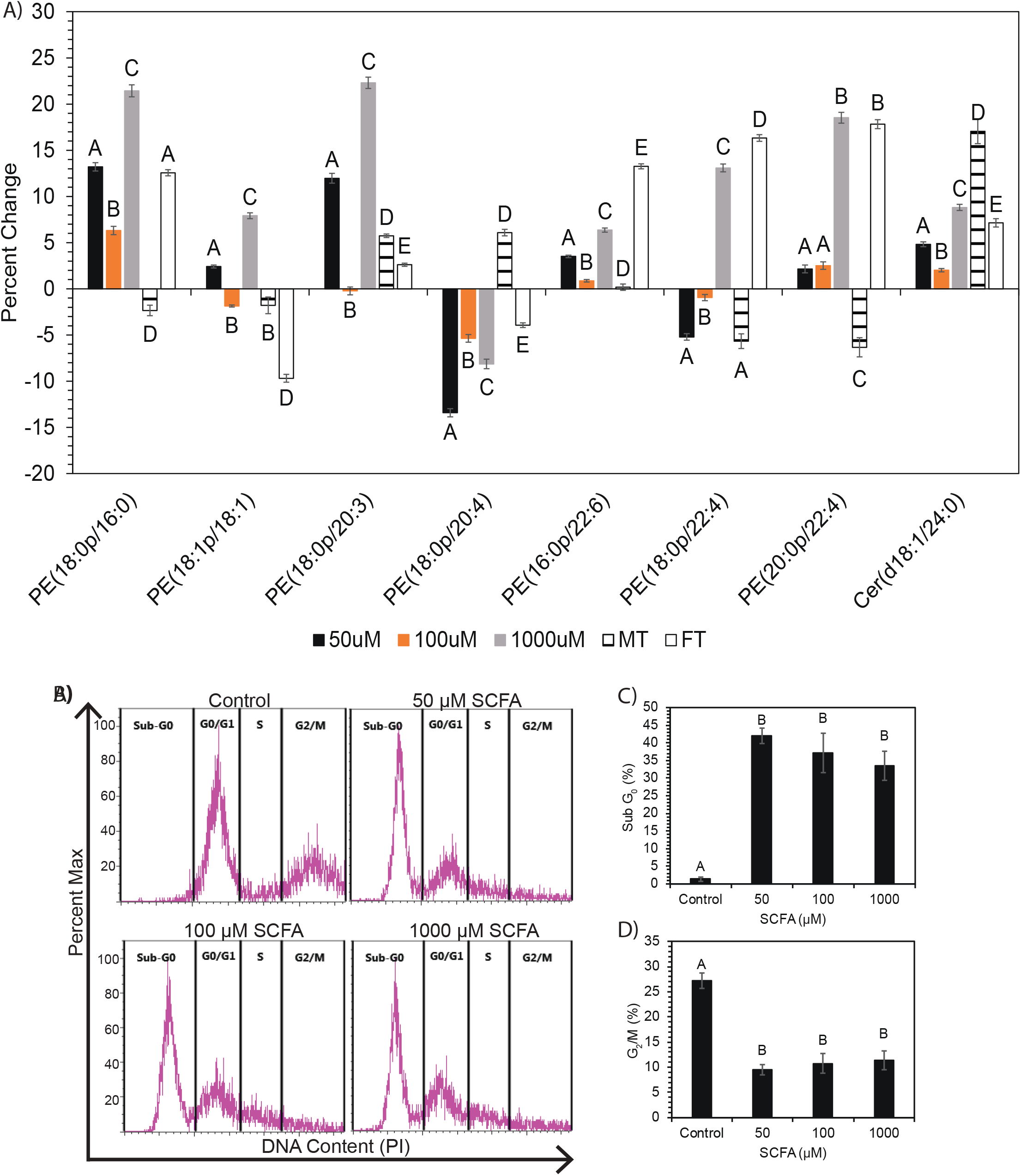
A) Percent change of treatments vs respective controls for ANOVA significant (p<0.05) plasmalogen and ceramide species altered in both SH-SY5Y cell line and Long Evans rat brain. Significance was determined via ANOVA following PCA analysis for plasmalogen and ceramide in-vitro and in-vivo. Bar charts representitive of mean ± standard error. Means represented by different superscripts are significantly different at p<0.05. MT = male rats treated with SCFAs. FT = female rats treated with SCFAs. (B-D) SCFA treatment of SH-SY5Y cells causes increased apoptosis accompanied by loss of cells in G_2_/M. B) Representative flow cytometry plots of propidium iodide (PI) staining of cells treated with control or 50-1000 μM SCFA. C) Frequency of cells in sub-G_0_ (apoptotic) and D) G_2_/M phases of the cell cycle. Bar charts representitive of mean ± standard error. Means represented by different superscripts are significantly different at p<0.05.

## Discussion

The brain is the most lipid rich organ in the body and consist of 60 % lipids on a dry mass basis. Thus, the brain lipidome is an important mediator of brain health and functional outcome. In fact, alterations to the brain cell membrane lipid composition can have numerous brain health implications related to the onset and progression of several neurological disorders ^(2, 31)^. For example, decreased PC and PE have been observed in Alzheimer’s disease (AD) patients ^(31)^, consistent with lipid alterations present within the neuronal cells in our study (Fig. 1D). Disturbance in lipid metabolism can be a trigger for inflammatory reactions, protein changes, and cell synaptic pathology, potentially escalating into neurodegenerative disorders such as Parkinson’s disease (PD) or AD ^(2, 16, 31)^. Furthermore, mitochondrial membrane lipid composition is intimately linked with mitochondrial structure and function, important brain health risk factors implicated in neurological disorders ^(2, 22, 26, 48, 53)^. Lipid comprised mitochondrial membranes are necessary for important cell functions including fission and fusion. Fission and Fusion are needed for cell cycle division and reproduction, while the lipid membranes allow for reservoirs of energy storage ^(2)^, which are crucial to brain health and function. The biggest component of these mitochondrial membranes are the high concentrations of phospholipids such as PC, PE (both of which constitute approximately 80% of the total brain phospholipids), and mitochondrial specific CL (10-15% of total lipids present in the brain) ^(2, 43)^. The mitochondrial composition is highly conserved between all mammalian cells, including humans ^(2)^.

In our study, we observed significant alterations in all major lipid classes after treatment with SCFAs. The most notable changes occurred in mitochondria associated lipids (Fig. 2-3). Neuronal carnitine showed several fold changes in response to SCFA treatment in both SH-SY5Y cell line and Long-Evans rat brain (Fig. 2). Carnitine can be derived either from the diet or synthesized within the brain and consists of both free carnitine and acylated carnitine species ^(11, 19, 44)^. Carnitine allows long-chain fatty acids to be transported across the inner mitochondrial membrane to be oxidized by beta-oxidation for energy or integrated into structural lipids. Alterations in these lipids can have metabolic and energy implications in the brain, as the brain switches from glucose to fatty acid energy under stress ^(11, 19, 44)^ such as during diseases or gut dysbiosis. Carnitine has been shown to have potentially neuroprotective effects in ameliorating metabolic disturbances in the brain and nervous system during injury. However, abnormal AC concentrations have been associated with autism, specifically elevated levels of short chain AC and long chain fatty acids ^(11, 13, 44)^, which were observed in both the cell line and rat brain analysis of this study (Fig. 2). Overall, there was an increase in free carnitine (C0), and all AC combined (C2-C20) (Fig. 1D) in neurons, consistent with the increased mitochondrial mass observed (Fig. 4A-B) following treatment with SCFA. This finding helps to connect the SCFA induced alterations in brain lipid metabolism to the impaired mitochondrial morphology, and the proposed brain health deficiency pathway reported in this study (Fig. 7). The high concentration of free carnitine found in this lipid profile is normal as adult neurons typically contain 80% free carnitine ^(19)^. Interestingly, both increases and decreases in free carnitine have been observed in mitochondrial dysfunction and neurological disorders ^(11, 13)^. In our study, increased C0 was observed, along with alterations in other groups and individual AC species in both neuronal cells and rat brain samples (Fig. 2). Several studies suggest that alterations in carnitine can augment aging related mitochondria, lipid, and metabolic dysfunction by impairing energy production and metabolism ^(11, 13, 19, 44)^. This supports carnitine’s ability to modulate brain energetics and its importance in maintaining overall brain health ^(19)^.

**Figure 7.**
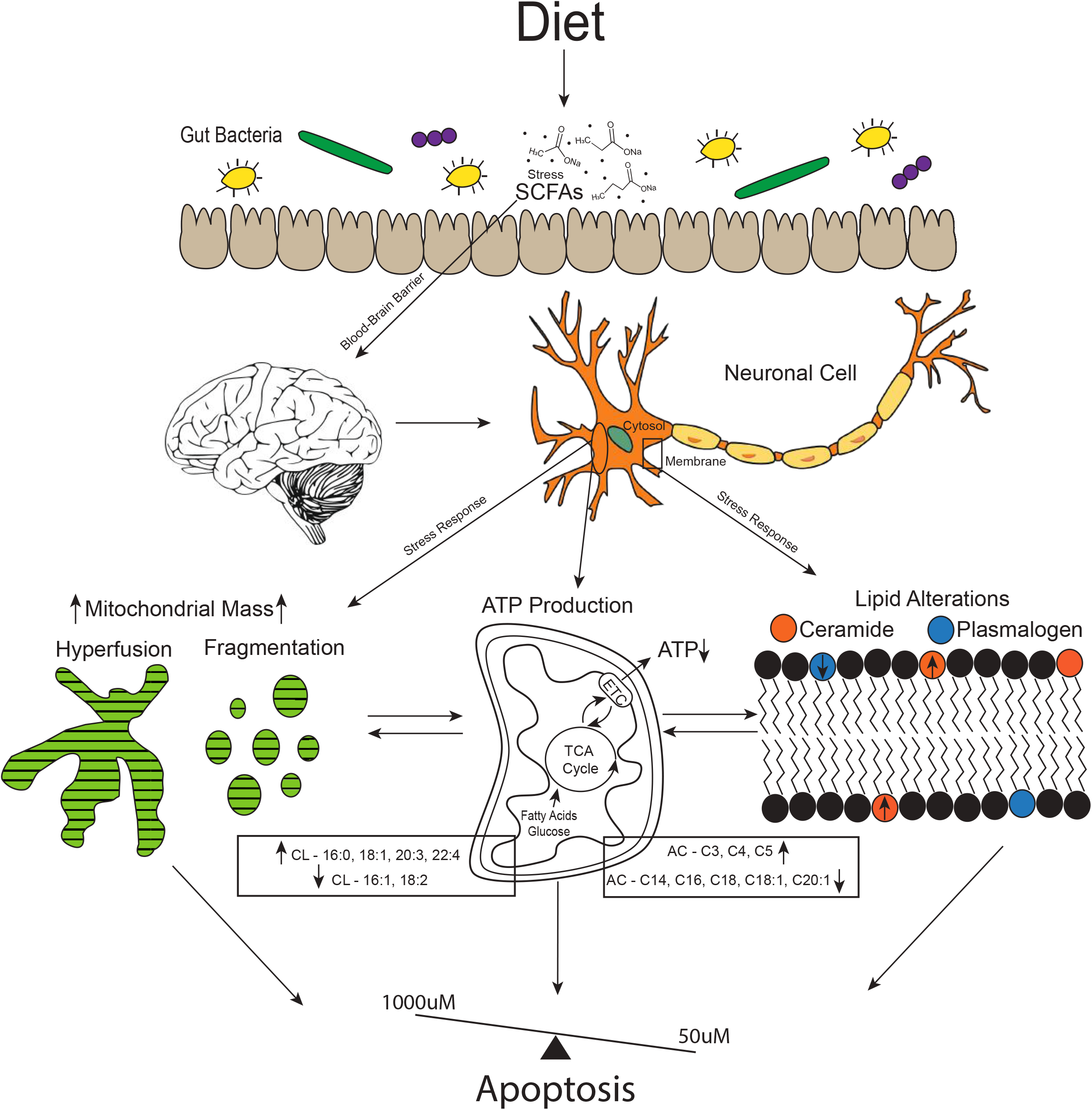
Proposed mechanism for neuronal cell response to levels (50-1000μM) of SCFAs observed in systemic circulation. SCFAs are recieved via infermentable carbohydrates in the diet, which are fermented by the gut microbiota population. SCFAs are able to cross the blood-brain barrier (BBB), which at elevated levels caused stress responses in brain neuronal cells. This stress response is shown in our study via alterations in lipids that make up the cell membrane bilayer, such as the increase observed in ceramide vs plasmalogen species. Mitochondrial lipid membrane is also altered, as observed by cardiolipin (CL) and acylcarnitine (AC) profiles. Mitochondria also display increased mass, hyperfusion (1000μM), fragmentation (50μM), and decreased ATP production. All of which are linked to apoptotic cell resonse, which we observed during cell cycle analysis. Although the 100μM treatment is not completely normal, it does not show major alterations compared to the 50μM and 1000μM treatments. This suggests a possible dose balance response between the treatments.

In this study, we also observed that SCFAs significantly altered the CL composition in neuronal cells (SH-SY5Y) and Long-Evans rat brain samples (Fig. 3). Decreased concentrations were seen among species dominated by 16:1 and 18:2 fatty acids, whereas increases were seen among species with 16:0, 18:1, and other long chain unsaturated fatty acids such as C20:3, C22:1, and C22:4. An overall increasing trend was observed in both neuronal cells and male treated Long Evans rats (Fig. 2B-E). CL is a mitochondrial specific lipid and can be an important biomarker to assess mitochondria function in neurons. Furthermore, CL also accounts for approximately 20% of the lipids in the mitochondrial membrane. In fact, CL is found almost exclusively in the inner mitochondrial membrane. CL plays a major role in ATP production and mitochondrial bioenergetics, as well as regulation of cytochrome c and apoptotic pathways. All these processes are essential for brain cell health and energetics, as disruptions in these processes can influence mitochondrial dysfunction and neuropathology ^(4, 29, 32, 34, 37)^. Alterations in CL are generally associated with pathologies such as mitochondrial dysfunction, oxidative stress, cell death, and/or aging ^(4, 32, 34, 37)^. Specific changes in composition, especially regarding aging, have not been commonly researched outside of the heart or cardiovascular system ^(4, 32, 34)^. However, within the aged heart, a decrease in 18:2 and an increase in other long chain polyunsaturated species have been observed ^(4)^. This is consistent with the decrease in 18:2 (Fig. 2B) and increase in 20:3, 22:1, and 22:4 species (Fig. 2C) observed in our cell line study, and the overall increase in polyunsaturated fatty acid (PUFA) enriched CL in the brain when male Long Evans rats are treated with SCFAs (Fig. 2E, Supplemental Data, Fig. 1). The alterations in both Car/AC and CL have clear connections to the observed alterations in mitochondrial morphology, ATP production, and cell respiration. These changes in CL are suspected to be connected to AD and/or PD, which may be induced by ATP changes and neuronal apoptosis ^(4, 29, 37)^. These finding suggests that exposure of neuronal cells to the elevated levels of SCFAs observed in patients during different pathologic and gut dysbiotic conditions appears to lead to an apoptotic response pathway that could possibly impact overall brain health over time (Fig. 7).

Furthermore, an increase in neuronal mass was observed (Fig. 4A-B), which correlated with increased long chain polyunsaturated CL species such as 20:3 and 22:4. Increased mitochondrial mass (high mitochondrial biogenesis) can be due to either increased energy demands or energy deficits such as cell stress, decreased ATP synthesis, or higher levels of oxidative stress, etc. ^(26, 45)^. This can be triggered by an increased demand for cellular energy that is unable to be met or by impairment of ATP production via the tricarboxylic acid (TCA) cycle ^(26, 45)^. In our study, we observe a decrease in ATP production in neuronal cells (Fig. 4F) and is congruent with the observed increased mitochondrial mass. The mechanisms behind increased mitochondrial mass are not well understood, but it may indicate an early event to prepare cells from oxidative stress via cell cycle arrest ^(26)^, which was also observed in this study (Fig. 6B-D). Abnormal mitochondria have been found under similar conditions in brain aging and neurological disorders ^(10, 26, 45)^. Mitochondrial mass has a positive correlation with CL content, where CL has been employed to represent mitochondrial mass ^(4)^. Based on the observed results, there is a clear connection between the SCFA induced alterations in brain mitochondrial lipid metabolism, morphology, and the neuronal cells apoptotic response (Fig. 7).

Furthermore, mitochondrial morphological analysis revealed high levels of mitochondrial fragmentation (50μM) and hyperfusion (1000μM) after treatment with SCFAs (Fig. 4C-D). Fragmentation is characterized by many small and round mitochondria, whereas hyperfusion is characterized by connected and elongated mitochondria ^(42)^. Fragmentation and hyperfusion play a role in cellular quality control, and cells continually alter the rate at which these occur based on energy demands ^(22, 38, 51)^. However, mitochondrial fragmentation is a hallmark of apoptosis and may result from nutrient overload (ie. SCFA). Furthermore, mitochondrial fragmentation can occur directly from apoptotic signals, or cause apoptotic signals ^(38, 45, 51, 52)^. While fragmentation is a normal and beneficial process in some functions of the body such as the increased energy demand of exercise ^(5)^, it has been observed in mitochondrial dysfunction related to overall brain health and aging, such as in PD and AD ^(47, 49, 52)^. We hypothesis that hyperfusion within the 1000μM treatment is likely a stress response to damaged molecules or mutated DNA, which at high levels or for long periods of time can lead to mitochondrial dysfunction ^(22, 38, 51)^.

Furthermore, although the 100μM treated cells display less than significant morphological changes (Fig. 4C-D), a reduction in ATP production and increased cell death was observed in this treatment concentration. These findings suggest that the increased mitochondrial mass and fragmentation/hyperfusion observed is a stress response affecting ATP production, further implicating the connection between mitochondrial mass and neuronal cell death by apoptosis following exposure to SCFAs (Fig. 7).

Based on mitochondrial lipid and morphology alterations connected to apoptosis (Fig. 7), we measured cell OCR via Seahorse XFe96 Mito Stress Test in live cells to further determine potential mitochondrial (dys)function. We observed a significant overall decrease in respiration (basal, maximal, spare respiratory capacity) and ATP production across all SCFA treatment levels (Fig. 4F). Brain cell energy metabolism is driven by mitochondrial respiration and ATP production within the mitochondria via oxidative phosphorylation, possibly affecting overall brain health. ATP is known to play an important role in oxidative stress, maintaining DNA, gene expression, and cognitive brain function ^(20)^. This is concurrent with our findings that 50μM treated cells were in a fragmented state after SCFA treatment, as ATP has been shown to stimulate fragmentation if decreased ^(22)^. When decreased, ATP productivity is also associated with mitochondrial dysfunction and oxidative stress, relating directly to diminished brain health in the form of neuronal damage, and aging related disorders, such as AD ^(20)^. Mitochondrial stress may lead to mitochondrial dysfunction if prolonged, potentially developing into neuronal cell death via apoptosis (Fig. 7). This reduction in respiration capacity and ATP production, coupled with subsequent lipid and morphological alterations, suggests overall impairment of brain health. Thus, marking SCFAs as a potential cause for mitochondrial stress and diminished brain health. ^(15, 20, 22)^.

It has been noted that all observed alterations to mitochondrial lipids, mitochondrial morphology, ATP, and cell respiration are connected via the potential for an apoptotic outcome due to stress inducing levels of SCFAs. Based on this information, we analyzed the apoptotic related lipids, plasmalogens and ceramides (Fig. 5-6A), as well as cell cycle progression (Fig. 6B). We observed a significant increase in the ratio of ceramides:plasmalogens, as well as a significant increase in Sub-G0 phase (apoptotic) cells and a significant decrease in G2/M phase cells. Recently, reduced plasmalogens were proposed as a risk factor for AD, which may occur simultaneously to increased ceramides ^(15, 18)^. Plasmalogens are ether phospholipids found mostly in PE or PC and play a major role in neuronal cell membrane composition ^(2, 15)^ and cellular stress response ^(8, 18)^. Furthermore, considering ceramides are suggested to be pro-apoptotic and plasmalogens anti-apoptotic ^(8, 15)^, the observed ceramide:plasmalogen ratio increase, due a higher fold-change by 15% in ceramides versus plasmalogens (Fig. 5A,C-D), suggests that a functional consequence of the altered mitochondrial morphology and lipid metabolism may be cell cycle arrest and eventual cell death via apoptosis. This is further supported by similar trends observed in rat brain plasmalogens. Specifically, the similarity between 1000μM treated neuronal cells and female treated rats (Fig. 6A). This shift shows that ceramide concentrations are more significantly increased compared to plasmalogen concentrations after SCFA treatment in-vitro and in-vivo; and this occurred concomitant with a significant decrease of neuronal cells present in the G2/M phase and increase in the Sub-G0 phase (apoptotic cells) in all SCFA treatment concentrations (Fig. 6B-D).

In this study, many of our findings point towards apoptosis, including lipid alterations, mitochondrial dysfunction via fragmentation/hyperfusion, respiration and ATP decline. These results present convincing evidence that SCFAs act as a stressor to neuronal cell health and viability and have major implications in neuronal cell survival by promoting alterations in lipid metabolism and mitochondrial morphology associated with an apoptotic response (Fig. 7).

## Conclusion

This study demonstrates that the elevated levels of SCFAs reported in systemic circulation of patience during gut dysbiosis and other pathological conditions significantly altered neuronal cells and Long-Evans rat brain lipidome, with the most noted alterations observed in mitochondrial specific lipids and PE plasmalogens. The altered mitochondrial lipidome occurred concomitant with hyperfused and fragmented mitochondria, decreased respiration, ATP production, impairment in neurons cell cycle progression, ultimately resulting in increased neuronal cell death by apoptosis. It is currently unclear as to whether the altered brain lipid metabolism, mitochondrial morphology, decreased ATP and respiration, or a combination of these are the trigger for the increased neuronal cell death by apoptosis observed following exposure to SCFAs. Based on the findings in this study, we propose a possible pathway by which elevated SCFA levels in the brain may translate into diminished brain health and neuronal cell viability. Our hope is that this work will stimulate further studies by the scientific community to elucidate how gut derived SCFAs modulate overall brain health outcome and function. We believe that the observed neuronal cells and equivalent rat brain responses to elevated SCFA levels reported in the systemic circulation are an important addition to the current literature and will be a seminal work leading to better understanding of how the gut microbiome modulate brain health outcome. Ongoing work in our research group is now focused on assessing the implication of the altered neurolipidome associations with any observed impaired behavioral phenotype and possible influence on brain health.

## List of Abbreviations

CNS: Central Nervous System
SCFAs: Short Chain Fatty Acids
C2: Acetate
C3: Propionate
C4: Butyrate
HDAC: Histone Deacetylase
HFD: High-fat Diets
IBDs: Irritable Bowel Diseases
CRC: Colorectal Cancers
FMF: Familial Mediterranean Fever
ROS: Reactive Oxygen Species
ATP: Adenosine Triphosphate
FBS: Fetal Bovine Serum
Pen/Strep: Penicillin/Streptomycin
PBS: Phosphate Buffered Saline
IP: Intraperitoneal
HILIC: Hydrophilic Interaction Liquid Chromatography
C30RPLC: C30 Reverse Phase Liquid Chromatography
PI: Propidium Iodide
PFA: Paraformaldehyde
MAP2: Microtubule-associated Protein 2
STED: Stimulated Emission Depletion
OCR: Oxygen Consumption Rate
FCCP: Carbonyl cyanide 4-(trifluoromethoxy)phenylhydrazone
XF: extracellular flux
PCA: Principle Components Analysis
ANOVA: Analysis of variance
LSD: Least Significant Difference
Cer & CerG1: Ceramides
LPE: Lyso-Phosphatidylethanolamine
CL: Cardiolipin
Car/C0: Carnitine
AC: Acylcarnitines
PE: Phosphatidylethanolamine
MT: Male treated
FT: Female treated
SRC: Spare Respiratory Capacity
AD: Alzheimer’s disease
PD: Parkinson’s disease
PUFA: Polyunsaturated Fatty Acid
TCA: Tricarboxylic Acid

## Declarations

### Ethics Approval

Outlined in Methods

### Consent for Publication

All authors consent for publication of this article

### Availability of Data

Not yet published but can be made readily available

### Author Contributions

T.A.F. wrote manuscript draft, R.H.T and K.M.D revised the final version. S.S., T.H.P., I.A., and M.L.T. contributed to data acquisition and interpretation.

S.K.C., and J.B. are co-supervisors on the project and have provided revisions. R.H.T. is the principal investigator (P.I.). All authors contributed to manuscript revision, read, and approved the submitted version.

## Acknowledgements

We acknowledge funding from Memorial University of Newfoundland (MUN) Multidisciplinary Fund, Natural Sciences and Engineering Research Council of Canada (NSERC), and the Aging Research Center Newfoundland and Labrador (ARCNL). Authors thank Dr. Christian Wurm of Abberior Instruments and his team for STED data acquisition and interpretation, as well as Dr. Tao Yuan for maintaining the laboratory instruments.

