## Supplemental Data for "Brief exposure of neuronal cells to levels of short chain fatty acids observed in human systemic circulation impair the cell lipid metabolism resulting in associated cell death by apoptosis"

| Features | p-value | Significant | MC | MT | FC | FT |
|---|---|---|---|---|---|---|
| CL(20:4/18:1/18:1/22:6) | 0.018 | Yes | 3.728 (a) | 3.618 (a) | 3.567 (a) | 4.050 (b) |
| **CL(20:4/18:1/20:4/18:2) | 0.018 | Yes | 3.626 (b) | 3.893 (b) | 3.157 (a) | 3.803 (b) |
| CL(22:6/18:1/18:1/18:1) | 0.040 | Yes | 4.554 (a) | 4.610 (a) | 4.681 (a) | 4.946 (b) |
| **CL(22:1/20:4/20:4/18:0) | 0.040 | Yes | 4.003 (b) | 3.811 (ab) | 4.176 (b) | 3.518 (a) |
| CL(23:0/16:0/18:0/22:6) | 0.040 | Yes | 4.862 (b) | 5.031 (b) | 5.220 (b) | 4.331 (a) |
| CL(20:4/18:1/18:1/20:4) | 0.043 | Yes | 5.414 (a) | 5.704 (a) | 5.632 (a) | 6.123 (b) |

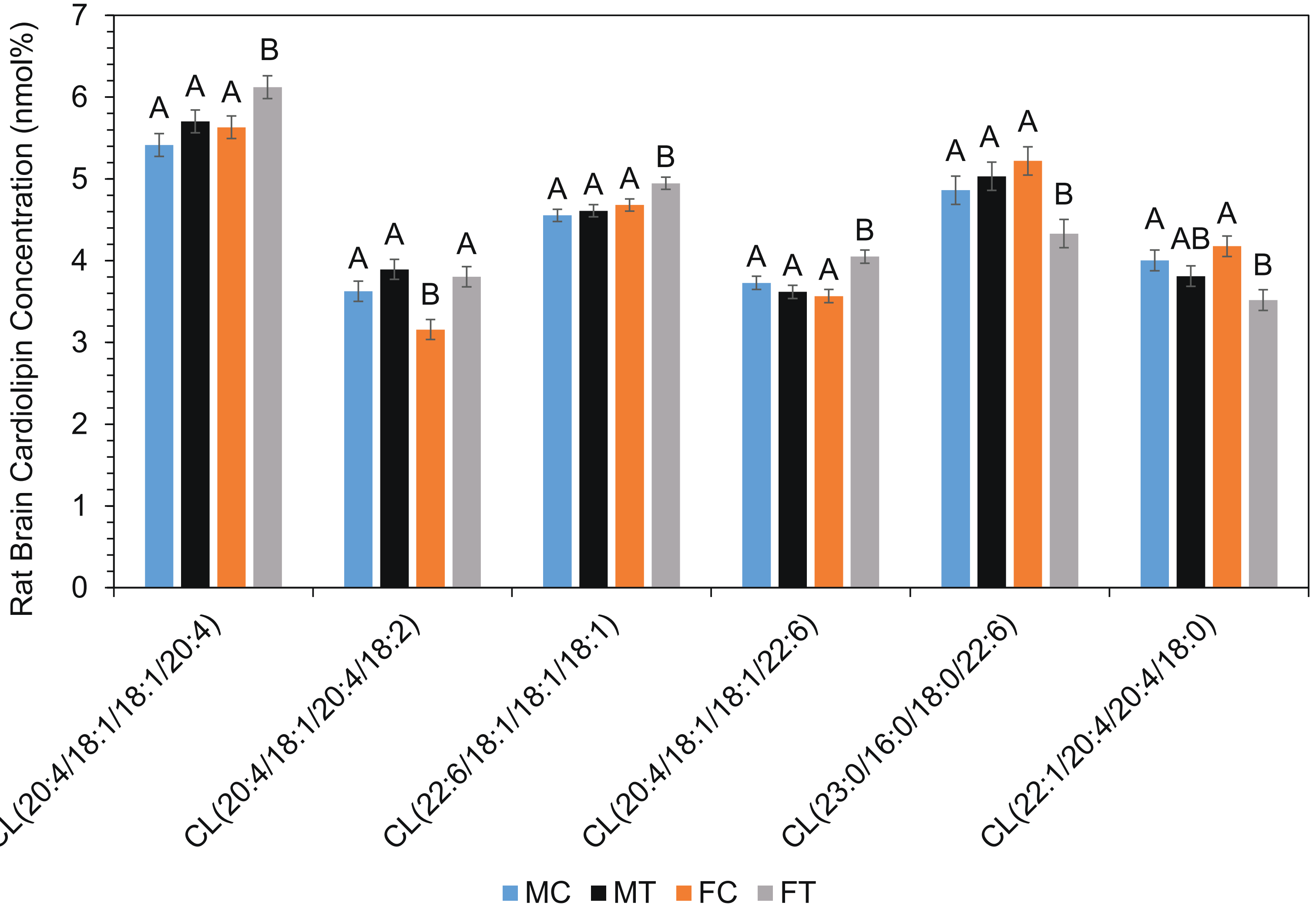


**Figure 1.** Alteration in cardiolipin (CL) composition after SCFA treatment. Differential Expression (A) was used to determine significant ($p<0.05$) alterations in molecular species between treatments. Significance in two-factor control versus treated analysis indicated by two asterisks - **. This result was also graphed (B). Bar charts representitive of means ± standard error. Means represented by different superscripts are significantly different at $p<0.05$.

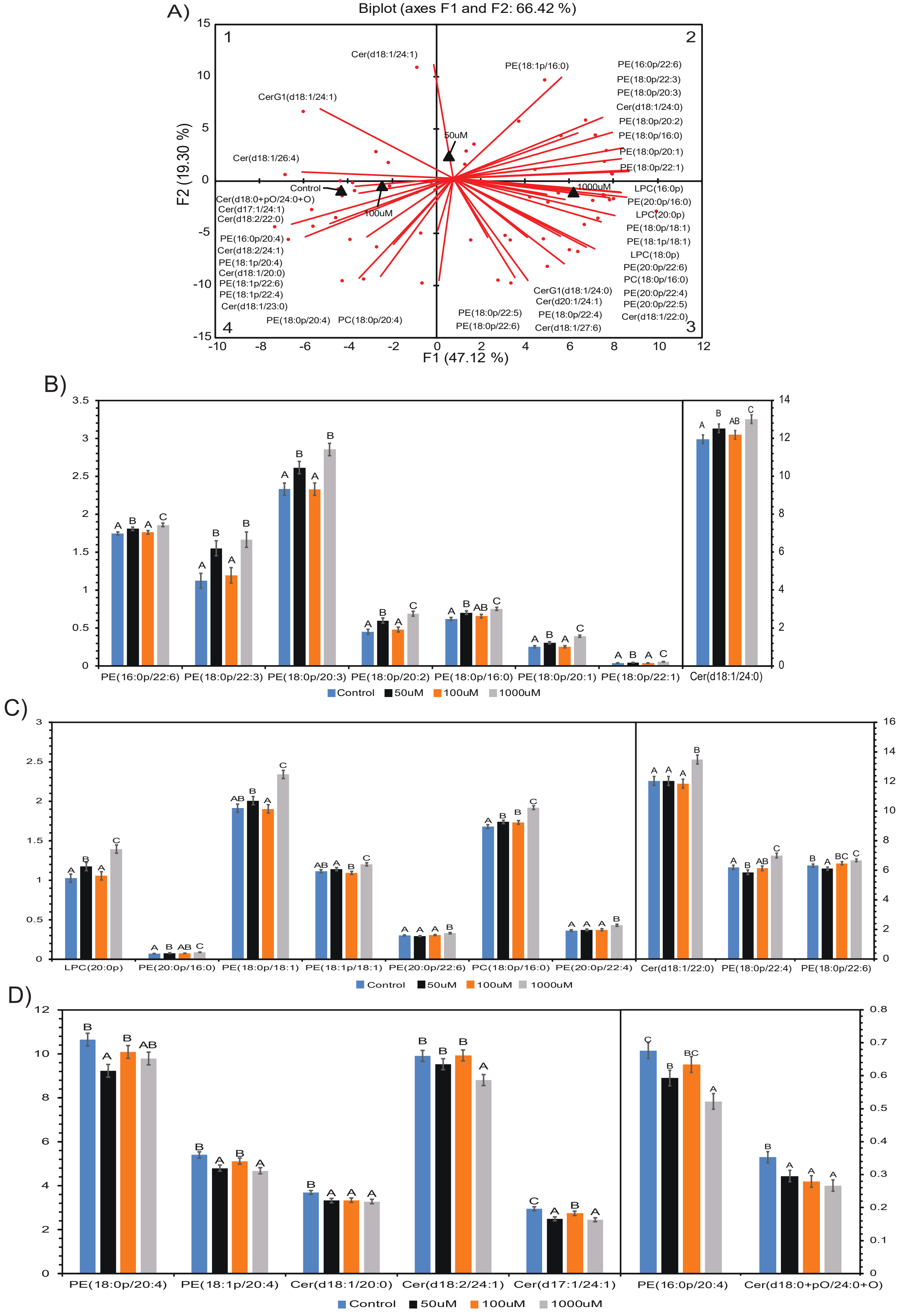


**Figure 2.** Alteration of plasmalogen and ceramide molecular species in neuronal cells. A) PCA biplot showing alteration in both ceramide and plasmalogen molecular species between control/100µM, 50µM, and 1000µM SCFA treatment. ANOVA was conducted based on PCA quadrants 2 (B), 3 (C), and 4 (D) corresponding to each treatment. Significant ($p<0.05$) molecular species are shown accordingly. Bar charts representitive of means ± standard error. Means represented by different superscripts are significantly different at $p<0.05$.
